# Sequence-based Prediction of Metamorphic Behavior in Proteins

**DOI:** 10.1101/2020.02.27.967935

**Authors:** N. Chen, M. Das, A. LiWang, L.-P. Wang

## Abstract

An increasing number of proteins have been demonstrated in recent years to adopt multiple three-dimensional folds with different functions. These metamorphic proteins are characterized by having two or more folds with significant differences in their secondary structure, where each fold is stabilized by a distinct local environment. So far, about 90 metamorphic proteins have been identified in the Protein Databank (PDB), but we and others hypothesize that a far greater number of metamorphic proteins remain undiscovered. In this work, we introduce a computational model to predict metamorphic behavior in proteins using only knowledge of the sequence. In this model, secondary structure prediction programs are used to calculate *diversity indices*, which are measures of uncertainty in predicted secondary structure at each position in the sequence; these are then used to assign protein sequences as likely to be metamorphic vs. monomorphic (i.e. having just one-fold). We constructed a reference dataset to train our classification method, which includes a novel compilation of 140 likely-monomorphic proteins, and a set of 201 metamorphic proteins taken from the literature. Our model is able to classify proteins as metamorphic vs. monomorphic with a Matthews correlation coefficient of ∼0.4 and true positive / true negative rates of ∼65% / 80%, suggesting that it is possible to predict metamorphic behavior in proteins using only sequence information.

**Statement of Significance:** This paper introduces the diversity index as a descriptor to distinguish metamorphic proteins, which possess multiple stable folds, from monomorphic proteins that possess only one fold. The diversity index is designed to measure uncertainty in computationally predicted secondary structure, which we hypothesize is elevated for metamorphic proteins. We tested our hypothesis by training a binary classifier using the diversity index and an annotated dataset of metamorphic and monomorphic proteins, and found an optimal Matthews correlation coefficient of 0.4, supporting the hypothesis and demonstrating for the first time that it is possible to predict metamorphic behavior in proteins using only sequence information. The sequence-based classifier has broader applicability compared to methods that rely on making comparison to experimentally measured structures.

## Introduction

Christian Anfinsen was awarded a Nobel Prize in Chemistry in 1972 for his work on the apparent one-to-one relationship between the amino acid sequence of a protein and its three-dimensional fold,^1,2^ giving rise to the classic paradigm: “one sequence, one fold”. However, serendipitous discoveries in the past few decades have led to the identification of “metamorphic proteins”^3^ that have the ability to jump reversibly between two distinctly different folds under native conditions. These proteins are fundamentally different^4^ from intrinsically disordered proteins^5^ (IDPs), morpheeins,^6^ and moonlight proteins^7,8^ which have been studied for a long time. Typical conformational changes in proteins often involve “shearing” or “hinge” behavior where entire protein subunits or secondary structure elements undergo relative motions without significantly altering the fold of the protein.^9,10^ In contrast, the different folds/structures of a metamorphic protein are dissimilar on a more fundamental level, often involving changes such as the transformation of a whole α-helix into a β-strand (Fig. 1). In this paper, we use *significant change in secondary structure* as the key defining characteristic of metamorphic proteins.

**Figure 1.**
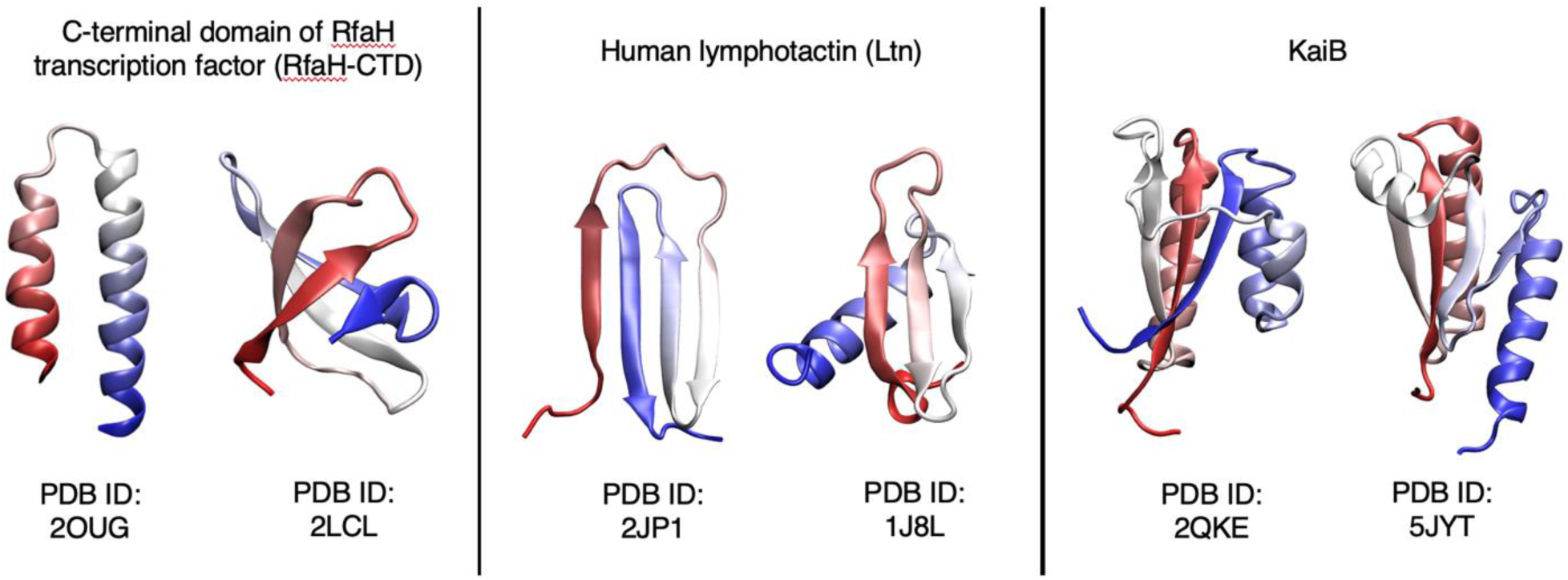
Representative examples of metamorphic proteins with 3-dimensional structures of both folds. The protein backbone is colored from N-terminal (red) to C-terminal (blue).

Although the number of known examples of metamorphic proteins such as IscU^11^, RfaH^12,13^, Selecase^14^, Mad2^15,16^, Lymphotactin^17^, CLIC1^18^, KaiB^19,20^ is relatively small, it is anticipated to increase steadily and populate the “Metamorphome”. In all these metamorphs, the transition from one-fold to another takes place in response to environmental triggers like pH, temperature, salt concentrations, binding partners, redox state, or oligomerization. Uncovering the metamorphome is crucial as it is expected have a transformative effect on long-held concepts of protein structure and function. It could also lead to engineering of metamorphic proteins, which are molecular switches, to act as sensors of small molecules or local environmental changes.

Traditional X-ray crystallography techniques, which account for solving 90% of the protein structures in the Protein Databank (PDB), are limited in their ability to identify metamorphism in proteins. These methods trap the protein in a minimum free energy structure in a specific crystallographic environment, thus they do not reveal the existence of alternate folds if the protein is metamorphic. A powerful method to detect protein metamorphism is solution-state NMR. However, high-throughput screening protein sequences for potential metamorphic behavior by NMR is not feasible. A realistic approach would be to identify metamorphic candidates using computational approaches, which would allow experimental verification to focus on a smaller set of candidate proteins.

A recent computational study from Porter and Looger^21^ identified 96 fold-switching candidates in the Protein Data Bank. The study stated that two characteristics of metamorphic proteins include discrepancies between experimentally derived and computationally predicted secondary structures, and the occurrence of multiple independent subdomains that each fold cooperatively. Using these two metrics, they estimated that up to 4% of the proteins in the PDB may be metamorphic, which suggests that this class of proteins appears to be more common than those identified so far.

In this work, we propose a novel binary classifier for predicting protein metamorphism based on the *diversity index*, which takes advantage of the uncertainty in secondary structure prediction methods. This method has a unique advantage that it can predict metamorphic behavior in a protein of interest purely based on the amino acid sequence, without requiring *a priori* experimental knowledge of the three-dimensional structure. The classification method is trained using two reference datasets consisting of ∼200 manually annotated monomorphic and metamorphic sequences respectively. We found robust performance of the diversity index-based classifier with a Matthews correlation coefficient of ∼0.4 (corresponding to ∼70% accuracy) that is largely insensitive to changes in the parameterization and training dataset.

The rest of this paper is organized as follows. We first give a brief overview of secondary structure prediction (SSP) methods, as they provide the essential inputs into our classifier. Next, we introduce the diversity index (DI) which measures the uncertainty of predicted secondary structure, and we outline how the DI is used to classify a protein sequence as metamorphic or monomorphic. This is followed by a description of the reference datasets containing known metamorphic and monomorphic sequences used to train our classifier. The performance of the classifier is discussed in detail using metrics such as the Matthews correlation coefficient, true positive rates, and true negative rates, and its robustness is tested using randomized cross-validation, sensitivity analysis, and examining how performance varies with different input SSP programs. We include a discussion of “outlier” protein sequences that are consistently misclassified by the DI-based model, as well as how the performance depends on the sequence database for position-specific scoring matrix generation, an important auxiliary input for the SSP programs. The paper concludes with some promising future directions.

## Theory

### Secondary Structure Prediction (SSP)

Secondary structure (SS) is a property of amino acid residues within a protein structure that describes its local intrachain three-dimensional structure. Under the well-known DSSP system,^22^ secondary structure may be classified into eight states, which can be further reduced down to three: α-helix (H), β-sheet (E), and random coil (C). In the years since the introduction of SS classification for known protein structure, several data-driven computational methods have emerged for secondary structure prediction (SSP) using only primary structure information, i.e. the amino acid sequence. Today, SSP is a vital part of the modern toolkit for protein structure prediction and design.

SSP methods can be understood in the conceptual framework of machine learning. The protein sequence is first processed into a feature vector consisting of information with structural relevance. Such features may include a PSSM (position-specific scoring matrix), which estimates the probability distribution of amino acid residues at each position in the sequence, and is computed by performing sequence alignments to a sequence database^23^ using programs such as PSI-BLAST^24^. The feature vector is input into a neutral network model, which has significant flexibility in its internal architecture, and provides three outputs representing the relative probabilities of helix (H), sheet (E), and random coil (C) at each position. The parameters of the neural network are trained to reproduce known secondary structures from widely available structural datasets. The accuracy of three-state SSP for modern methods has been reported to be as high as 82-84%.^25^

In this paper, four widely used SSP programs were applied to predict the secondary structure of every sequence in our datasets, namely Psipred^26^, SPIDER2^27^, SPIDER3^28^, and Porter 5.0, denoted here as Porter5^29^. Psipred, developed in 1999, introduced the idea of using the PSSM generated by PSI-BLAST as input to a neural network for secondary structure prediction. SPIDER2 uses a deep neural network that incorporates the PSSM from PSI-BLAST along with amino acid physicochemical properties^30^ to predict secondary structures and main chain dihedral angles. SPIDER3 is an updated version of SPIDER2 that incorporates hidden Markov model sequence profiles generated by the HHBlits program^31^ as input to a bidirectional recurrent neural network architecture, effectively allowing the entire sequence to calculate SS prediction at each position instead of a sliding window as in SPIDER2. Porter5 is the latest version of a series of SSP programs and uses HHBlits-generated HMM sequence profiles and PSI-BLAST-generated PSSMs as input. In this paper, we used the UniProt90_2019_01 sequence database as the input to PSI-BLAST for PSSM generation, and the Uniclust30_2018_08 database was used as input for HHBlits. These published sequence alignment databases are distinct from the metamorphic and monomorphic reference datasets that were compiled as part of this work.

### Metamorphic Proteins and Diversity Index

Metamorphic proteins can reversibly adopt multiple folded conformations for the same amino acid sequence under native conditions.^3,4^ Moreover, representative examples of metamorphic proteins are characterized by significant differences in secondary structure between folds (Figure 1), which is a distinct feature from more typical kinds of conformational change that generally preserve secondary structure as described in the Introduction. Because these metamorphic proteins possess multiple stable folds with differences in secondary structure, our central hypothesis is that metamorphic protein sequences are able to “confuse” secondary structure prediction programs. According to this hypothesis, we defined one descriptor, the diversity index (DI):

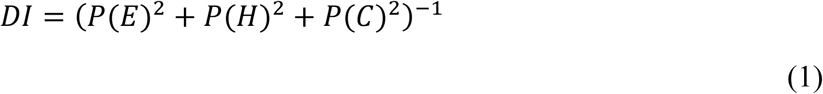

where *P(E), P(H)* and *P(C)* are output quantities from the SSP program representing the probabilities of strand, helix and coil, respectively, for a single residue in the sequence. The DI for a residue ranges from 1 to 3. When the value is close to 1, it means the SSP program has a high degree of confidence in the predicted SS, whereas the predicted SS is less certain when the DI value is close to 3. Because metamorphic proteins tend to have contiguous portions of the sequence (or even the whole protein) that undergo changes in secondary structure, we also hypothesized that the DI of metamorphic protein sequences will be elevated in contiguous regions of the sequence. Therefore, we consider the maximum value of a moving average of the DI over the sequence as the main criterion to predict metamorphic behavior in a protein sequence. In other words, a sequence will be classified as metamorphic if the following criterion is satisfied:

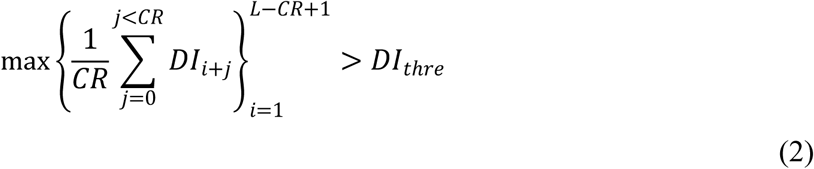

where the *CR* “number of consecutive residues” and *DI*_*thre*_ “diversity index threshold” are adjustable parameters. This binary classifier needs to be trained on reference or “manually annotated” datasets consisting of known-metamorphic and known-monomorphic (i.e. single fold) sequences. We will describe the construction of these datasets in the following sections.

## Dataset Setup

### Construction of the metamorphic reference dataset

In 2018, Porter *et al* published a paper listing 192 (96 pairs) of existing metamorphic proteins^21^ in which most pairs have very high sequence similarity to one another (between 90% and 100%). Our metamorphic reference dataset, listed in Supplementary Table S1, makes the following revisions to the listing in Ref. 21. In eight cases, both folds of a metamorphic protein existed as different chains with identical sequences in a single structure, and these sequences are counted twice in our dataset (e.g. 5C1V). Among the original set of structures, one protein is no longer available from the database (PDB ID: 2A01). We also removed proteins where the fold switching region is contained within 20 residues of the N- and C-termini (4ZRB, counted twice) or if the sequence length is shorter than 40 residues (4FU4, 4G0D, 5K5G and 2KB8); this is because our classifier requires taking a moving average of the diversity index, requiring an sequence that is longer than the largest window size (>15 residues) plus the number of the removed terminal residues (>5*2 residues). In total, 8 proteins were removed from the list for the reasons above. We also added several proteins to the list, including the known metamorphic protein GABA that was excluded from Ref. 21 (PDB ID: 2LHC and 2LHD) and 15 other possible metamorphic proteins which only have one fold of solved structures but experimental evidence for metamorphism, such as 2LSH. Our reference metamorphic dataset contains 201 metamorphic proteins in total.

### Construction of the monomorphic reference dataset

Our classification model for predicting protein metamorphism needs to be trained on proteins with known metamorphic behavior, as well as those with known single-fold (i.e. monomorphic) behavior. Although it is widely assumed that the PDB contains mostly monomorphic proteins, it is likely that a significant portion exhibits as-yet undiscovered metamorphic behavior. Therefore, we queried the PDB to obtain a set of protein structures that are *highly likely* to be monomorphic based on the following set of criteria: (1) The structure should be reported at least 10 years ago and has a good quality structure, in the sense that X-ray structures with resolution > 2.2 Å were filtered out; (2) there must be >30 published structures with at least 50% sequence similarity with the structure of interest; (3) the sequence length is > 40 and < 250 residues, in order to meet the criteria of having a well-folded core while staying within the typical sequence lengths of globular proteins. Each structure found in the above manner is termed ‘parent protein’, and structures with high sequence similarity found in step (2) above are termed ‘child proteins’. A total of 1387 ‘parent protein’ structures with a maximum sequence similarity of 70% and more than 65000 ‘child protein’ structures were downloaded along with their abstracts from the RCSB PDB web server using an automated crawler written in Python that uses the *scrapy* package. Two filtering rules were imposed in order to maximize the probability that a ‘parent protein’ is monomorphic:

1. The root-mean-square deviation (RMSD) values were calculated for all pairs of structures after sequence alignment for a ‘parent protein’ and all of its ‘children’. The structure was excluded from the data set if any of the pairwise RMSD values exceeded 2.4 Å.
2. The mismatch in secondary structure (SS) was calculated for all pairs of structures after sequence alignment with a ‘parent protein’ and all of its ‘children’. A positional mismatch score is calculated by summation over aligned residues in a window of 30 residues in length, where “2” was assigned if one sequence is H and the other is E, and “1” was assigned if one sequence is C and the other is either E or H, then taking the maximum value over all window positions. The structure was excluded from the data set if the SS mismatch score between any pair of sequences exceeded 9.

Finally, the abstracts of the corresponding publications were checked for keywords such as ‘fold-switching’, ‘metamorphic’, ‘two-folds’, and other synonyms; if the abstract indicated possible metamorphic behavior, then it was excluded from this dataset as well. This procedure resulted in a total of 140 likely monomorphic proteins (Supplementary Table S2), including 4 proteins that we deemed to be monomorphic from reviewing the literature but did not meet the above criteria.

An example of a metamorphic protein (KaiB) and a monomorphic protein (1AB9) from our reference datasets is shown in Figure 2. The highest RMSD value in the KaiB (typical example of metamorphic proteins) cluster exceeds 7.0 Å, and many pairs of sequences exhibit a secondary structure mismatch of 23 or greater. On the contrary, both the RMSD values and SS score are consistently low for the monomorphic protein, 1AB9.

**Figure 2.**
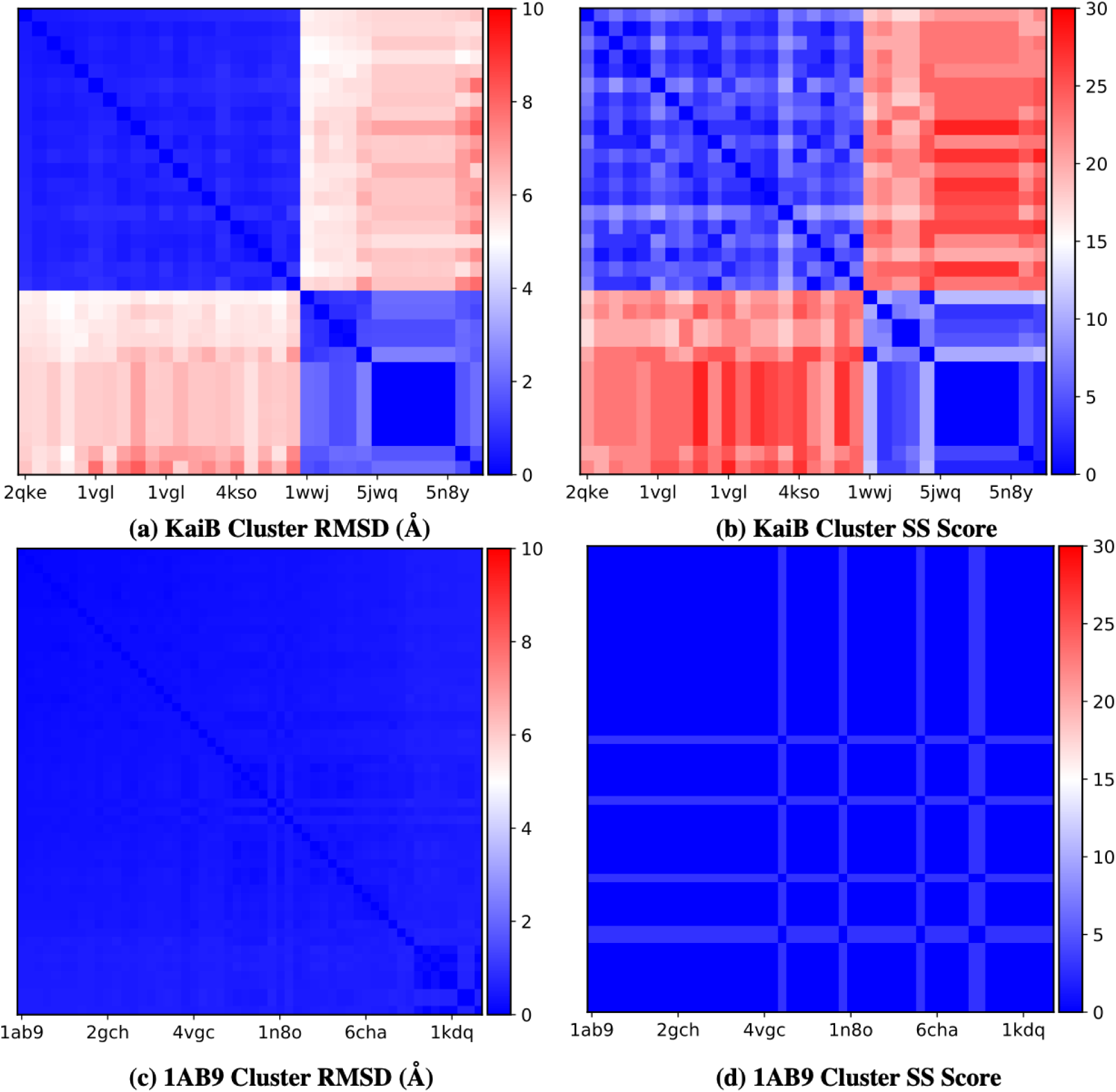
RMSD (a,c) and SS (b,d) score of KaiB and 1AB9, respectively. The highest RMSD value and highest SS score of the KaiB cluster is much larger than those of the 1AB9 cluster.

## Results and Discussion

### Behavior of the diversity index (DI)

According to Equation (1), the range of the diversity index (DI) is from 1 to 3, with larger values indicating greater uncertainty of SS prediction. Figure 3 plots the SS and DI from the SPIDER2 program for a well-known metamorphic sequence (KaiB, left panel) and monomorphic sequence (ubiquitin, right panel) along with the experimentally derived secondary structure(s). As shown in the left panel, the DI of the KaiB sequence has several regions of elevated values in the metamorphic region that spans positions 60-90. On the other hand, DI of ubiquitin is relatively low for the whole sequence, with small jumps at the boundaries of different secondary structure domains that are smoothed out by taking the moving average. This example illustrates how diversity indices may be used to predict metamorphic behavior in proteins when the folds exhibit different secondary structure in the metamorphic regions.

**Figure 3.**
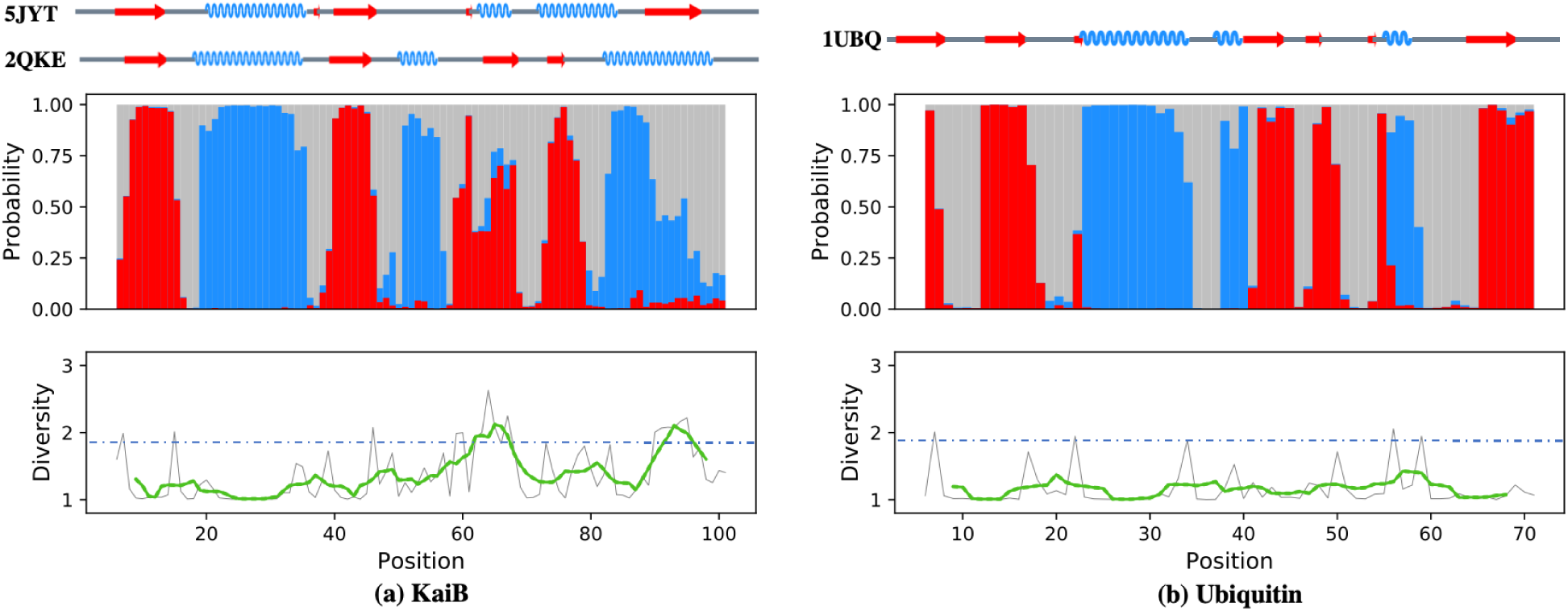
Secondary structure prediction results for KaiB (a) and Ubiquitin (b). Top: Sequences depict the experimentally derived SS of both KaiB structures (PDB ID: 2QKE and 5JYT, left) and Ubiquitin (1UBQ, right). Middle panel: Predicted SS probabilities at each position in the sequence from SPIDER2 (red, strand (E); blue, helix (H); gray, coil (C)). Bottom panel: The diversity indices for each residue position in the sequence (gray), with a moving average with a window size of 14 (green) and DI threshold for metamorphic behavior (blue dotted line). The DI takes on higher values when the predicted SS is more evenly distributed between H, C, and E, indicating greater uncertainty. For KaiB, the higher DI regions coincide with the experimentally known metamorphic regions.^19^

### Diversity index-based classifier performance

The performance of our model is measured using the Matthews correlation coefficient (MCC), a well-established measure of quality of binary classifications. For each combination of our parameters *CR* and *DI*_*thre*_, the MCC is computed from a matrix of true and false positives and negatives, called a confusion matrix.^32^ Generally speaking, larger values of *CR* correspond to increased window size and tend to decrease the maximum value of the moving average. Larger values of *DI*_*thre*_ also tend to decrease the probability that a sequence is classified as metamorphic. Thus, for increasing values of *CR* and *DI*_*thre*_, the true negative and false negative rates both increase. In order to get a better understanding into the behavior of our model, we plotted heat maps of the MCC in our two-dimensional parameter space. Herein, we consider possible values of *CR* ranging from 6 to 15, and possible values of *DI*_*thre*_ ranging from 1.4 to 2.6 with a step size of 0.05.

The sensitivity of our model was tested by cross-validation. In each of 10 trials, the reference metamorphic and monomorphic data sets were each randomly divided into training and test sets comprising 85% and 15% of the data, respectively. The parameters were determined by maximizing the MCC for the training set, then these parameter values were used to calculate the MCC for the test set.

Figure 4 and Table 1 show the main results for our DI-based classification using four SSP programs. Similar levels of performance for the training set were obtained using all four SSP programs as input to the DI-based classification. Among these methods, SPIDER2 had the largest average MCC value of 0.421 for the training set (Table 1), which was slightly higher than that of Porter5 (0.401), SPIDER3 (0.382) and Psipred (0.398); the differences were rather small and within the standard errors from randomized cross-validation trials. The parameters that maximized the MCC tended to appear in the middle of the parameter space, with significant regions of the parameter space exhibiting only minor variations from the optimum. For example, in the case of SPIDER2, the largest MCC value among all the trials was around *CR* ∼14 and *DI*_*thre*_ ∼2.1.

**Table 1.** The results from four different SSP programs, including the training set results and the test set results. The numbers in parentheses were the sample standard deviations.

| **MCC** | **Psipred** | **SPIDER2** | **SPIDER3** | **Porter5** |
| --- | --- | --- | --- | --- |
| Training Set | 0.398 (0.023) | **0.421 (0.015)** | 0.382 (0.015) | 0.401 (0.014) |
| Test Set | 0.315 (0.088) | 0.338 (0.084) | 0.286 (0.103) | 0.313 (0.083) |

**Figure 4.**
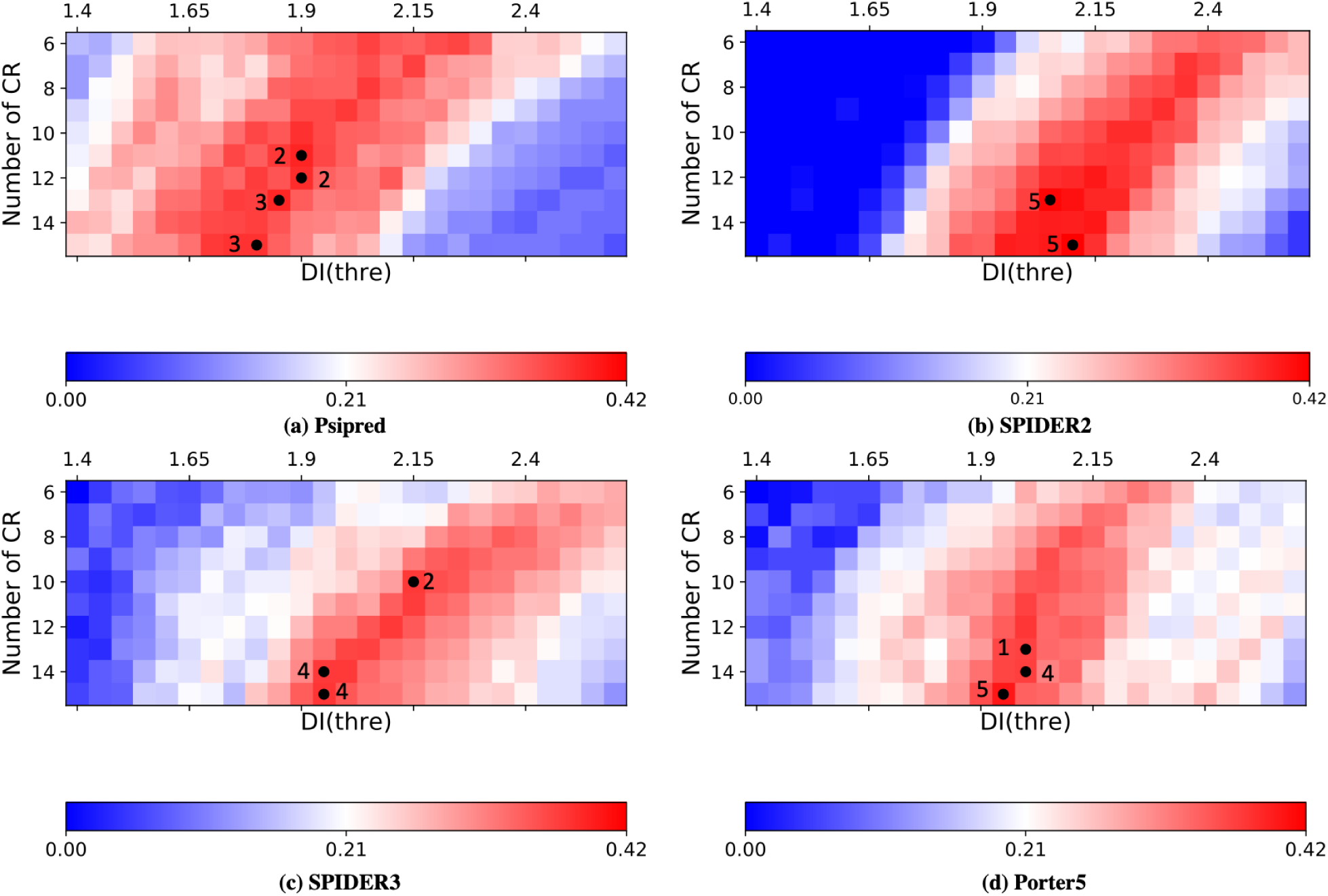
MCC heat maps for the diversity index-based classifier using predicted secondary structure from four programs, namely (a) Psipred, (b) SPIDER2, (c) SPIDER3 and (d) Porter5. The color map (blue < white < red) correspond to MCC values computed for the full reference dataset. Each point indicates the optimized parameter value for a randomly selected training set (85% of the full dataset), with numbers indicating how many times each optimum was found.

In terms of the test set, Porter5 and Psipred performed similarly with MCC values of 0.31–0.32, with differences being within the standard errors from randomized cross-validation trials. SPIDER2 has the highest average MCC value (0.338) for the test set. The small difference between the test set and training set MCCs, and the consistency of our results across several models, indicate that the DI-based classifier is a robust method for predicting metamorphic behavior. SPIDER3 showed lower performance in the test set compared with the other three methods, with a MCC of 0.282. Figure 4 also shows that SPIDER2 has a broader range of parameter space with near-optimal performance as compared to Psipred, SPIDER3 and Porter5. The training and test results overall indicate that higher secondary structure prediction accuracy does not directly translate to better performance in metamorphic protein classification.

Although these methods had similar MCC values, their accuracy in terms of correctly predicting true positives and true negatives showed much greater variations. According to the data shown in Table 2, the true positive rate (TPR) is lower than the true negative rate (TNR) in all four methods for the optimum parameters that maximized the MCC. Among these methods, SPIDER3 has by far the highest TNR value (0.92) and lowest TPR (0.42). The other three methods had similar TPR ranging from 0.59–0.66 and TNR ranging from 0.78–0.83, which are within the limits of statistical errors from our cross-validation studies. We presumed that the large TNR values of SPIDER3 comes from overall low values of the calculated diversity index, which possibly originates from higher SS prediction confidence levels as compared to other methods. We thus recommend SPIDER2 as the input method of choice for metamorphic protein classification, due to its consistently high MCC value for both training and test sets, balanced true positive and true negative rates, and wide regions of parameter space with near-optimal performance.

**Table 2.** The positive and negative accuracy of four different SSP programs

| **Accuracy** | **Psipred** | **SPIDER2** | **SPIDER3** | **Porter5** |
| --- | --- | --- | --- | --- |
| True Positive Rate | 0.590 (0.046) | 0.654 (0.038) | 0.416 (0.085) | 0.618 (0.018) |
| True Negative Rate | 0.828 (0.042) | 0.791 (0.041) | 0.917 (0.031) | 0.787 (0.017) |

### Comparison with other methods

There currently exist a few methods in the literature for predicting metamorphic behavior in proteins.^21,33^ To our knowledge, all existing methods require the knowledge of either the protein’s three-dimensional structure or secondary structure information from experiment. Porter *et al*. hypothesized that metamorphic proteins possess at least one independent folding domain with a 3-D structure that is largely independent of the rest of the sequence and proposed a method to predict metamorphic behavior based on prediction of independent folding domains.^21,34^ This method uses the protein’s 3D structure as essential input data, and thus its predictions are based on existing structural knowledge.

More recently, Porter *et al* and coworkers reported that metamorphic proteins have lower SSP accuracy than monomorphic proteins or fragments,^33^ which is similar to the ideas in our current work; however, the method they proposed requires prior knowledge of experimental secondary structure. A major differentiating feature of the diversity index-based classification method presented here is that it requires no experimental data for the sequence of interest. Thus, this method could be used to make predictions of metamorphism in protein sequences where there is no existing structural data.

### Classification using multiple diversity indices

We also examined the possibility of obtaining an improved classification model based on a linear combination of DIs obtained from two SSP programs, essentially increasing the number of descriptors to two. The discriminant parameters (i.e. slope and intercept of the line) were optimized by maximizing the MCC.^35^ Using a linear combination of the SPIDER2 DI and the Porter5 DI, we found the MCC value of the optimal model increases to 0.45. Figure 5 plots the discriminant line and the descriptor values for each protein as a scatter plot. The diagonal shape of the distribution indicates a high degree of correlation between the two diversity indices (R^2^=0.41), and most of the metamorphic proteins identified as true positives (TP) are located in the top-right corner of the figure. We found similar performance using some alternate approaches, for example an “inconsistency index” to predict metamorphism using the level of disagreement between two SSP programs (Supplementary Figure S1), and principal component analysis on the results of multiple SSP programs followed by K-means clustering (Supplementary Figure S2). These methods all yielded results with MCC values within 0.1 of the basic method using a single diversity index.

**Figure 5.**
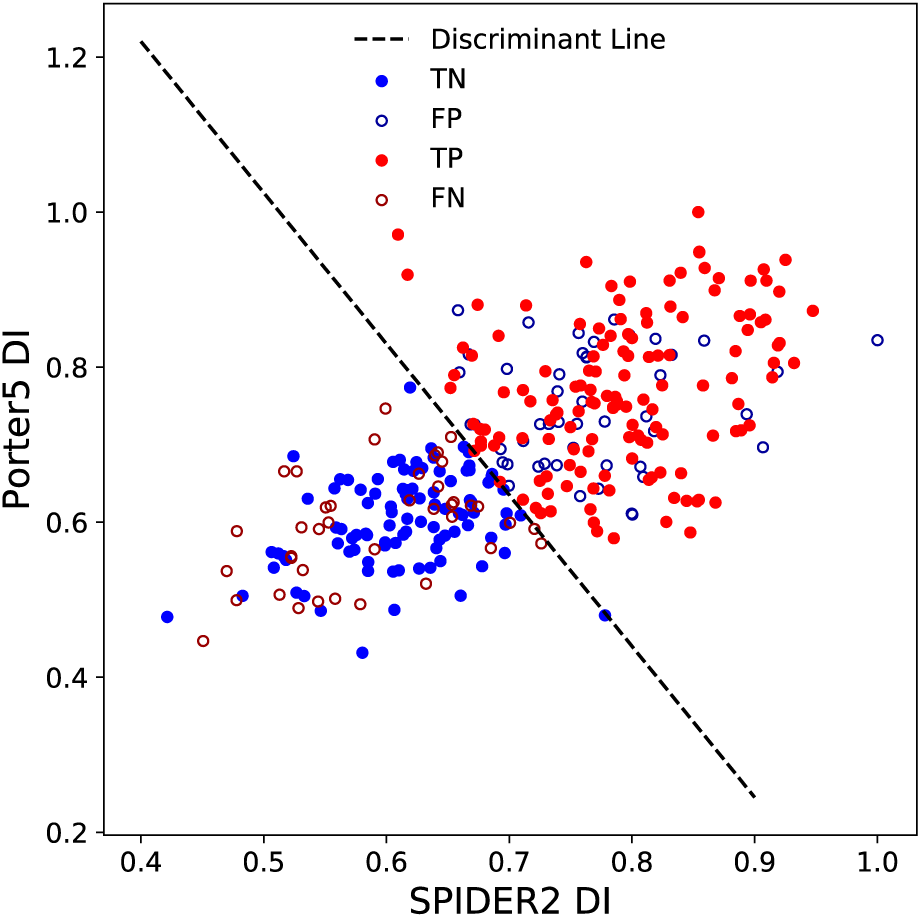
Two-parameter classification using diversity indices from SPIDER2 and Porter5. Here the true positives (metamorphic proteins in the reference dataset correctly classified as metamorphic) are represented by red filled circles, and the false negatives (metamorphic proteins in the reference dataset incorrectly classified as monomorphic) are represented by red open circles. The blue filled circles and blue open circles represent for true negatives (monomorphic proteins in the reference dataset classified as monomorphic) and false positives (monomorphic proteins in the reference dataset classified as monomorphic), respectively.

However, our analysis also revealed some false negatives (FN, open red circles) in the lower left of the Figure 5; these are metamorphic proteins in our reference dataset but have very low diversity indices, and contradict our rationale for the DI-based classifier. The same applies for false positives (FP, open blue circles) in the upper right of the Figure 5, as these are monomorphic proteins in the reference dataset with high diversity indices. In the following section, we provide a rationale to explain these outliers.

### Analysis of outliers in diversity index-based classification

Several proteins are consistently misclassified by the DI method using the predicted SS from all four programs. There are 22 metamorphic proteins in our reference dataset that are consistently misclassified as monomorphic proteins (false negatives), and 14 monomorphic proteins consistently misclassified as metamorphic proteins (false positives).

We examined the 22 “persistent” false negatives, i.e. metamorphic proteins from our reference dataset that are consistently mis-classified as monomorphic, and generally found that their two folds did not satisfy our initial criterion of having significantly different secondary structures, and instead feature other kinds of conformational differences, which we discuss in the following examples. The three-dimensional structures of the false negatives are shown in Supplementary Figure S3.

1. 2LQW^36^:2BZY^37^. A closer examination reveals that these two structures have very similar secondary structures. As shown in Figure S2, 2LQW is a key signaling protein that exists as a monomer while 2BZY is a partial structure of a CrkL homo-dimer protein, in which the existing part has similar SS to 2LQW. Moreover, the truncated CrkL monomer protein (PDB ID 2BZX) has highly similar secondary structure to 2BZY. The high similarity in secondary structure is consistent with the classification assigned by our method, which is based on differences in secondary structure between folds.
2. 2NNT^38^:2MWF^39^. 2NNT is a tetramer amyloid protofilament that forms an extended β-sheet between multiple chains, whereas 2MWF is a mutant monomer that forms a β-sheet within the residues in one chain. Again, the highly similar secondary structure in both folds is consistent with the classification assigned by our method.
3. 4HDD^40^:2LEP^41^. The structures in this pair are similar in terms of secondary structure but have a large RMSD. 4HDD is a homodimer in which a β-sheet is formed between chains, whereas 2LEP uses the same domain to form a β-sheet within one chain.
4. 1G2C^42^ is a truncated protein whose SS closely matches with the corresponding residues in 5C6B, which is a full structure. Strikingly, the other protein 5C6B^36^, which has similar sequence to 1G2C, is correctly classified by SPIDER2 and PORTER5 as a metamorphic protein. The high-DI domain of 5C6B (residue 270 to 295) was not part of the 1G2C structure, which indicates the incorrect classification of 1G2C might be solely due to the truncation of the input sequence.
5. 4XWS^43^:4Y0M^44^. Upon examination of the structures, we think this structure pair had been incorrectly included in our reference metamorphic dataset, as these two structures are highly similar in terms of secondary structure as well as three-dimensional structure (RMSD: 1.561 Å). In fact, the text of Ref. ^43^ states that the metamorphic region of the protein could not be solved by X-ray crystallography.

Within the 14 persistent false positives, i.e. monomorphic proteins from our reference dataset that are consistently classified as metamorphic, we found the following examples, with 3D structures shown in Supplementary Figure S4:

1. 2UU8^45^ is a concanavalin A protein and its DI value is relatively high in all the SSP programs, particularly in SPIDER2 (about 2.5). This structure possesses many short adjacent domains with different secondary structure, which leads to uncertainty in the SSP programs. Also, β-sheets are dominant in the SS of 2UU8, and we observed that the outermost strand of the β-sheet has a general tendency to have high uncertainty from SSP programs.
2. 3SEB^46^ is a protein that with several short SS domains (including α-helices and β-sheets), leading to the high DI values for these domains, similar to the example above. More than half of the false positives follow the same trend, indicating that our DI-based classifier is biased to misclassify protein sequences that are monomorphic but intrinsically difficult for SSP programs due to having many short subdomains with distinct SS or outermost β-sheet among several (anti-)parallel β-sheet strands.
3. 2JE7^47^, a recombinant lectin, has the same situation that the outermost strand of β-sheet and the short adjacent SS domains have the highest DI values. However, this protein is known to form either a dimer or a tetramer depending on the pH value. Although no direct evidence shows the SS change during this dimer-tetramer equilibrium process, it is possible that this process is associated with metamorphism not yet discovered.
4. 3CHB is another labeled monomorphic protein whose DIs are very large in all the SSP programs. Unlike the other two false positive proteins above, 3CHB^48^ has a long α-helix and five medium length β-sheets. Another short length α-helix is located at the N-terminal. According to the SPIDER2 prediction, three out of five β-sheets have relatively large DIs, resulting to a region with high average DI value. So far, we do not have a good explanation for the reason of this false positive. One possibility is that other proteins in the PDB have highly similar sequences to these high-DI β-sheets but have different SS.

We note that it is possible for our reference monomorphic dataset to include proteins that are actually metamorphic, despite our efforts to minimize this occurrence. This is because our selection of monomorphic proteins was based on the analysis of known structures in the PDB, which by definition excludes alternate folds or structures that have not yet been discovered or deposited.

### Dependence of results on sequence database

The performance of SSP programs relies on the non-redundant sequence database that is used to compute the position-specific scoring matrix. Figure 6 shows the differences in classification performance when SPIDER2 is used as the SSP program for different choices of the non-redundant sequence database. The Uniref-50, Uniref-90 and Uniref-100 databases have a progressively larger number of sequences and sequence identity among pairs of sequences. Figure 6 shows that Uniref-50 has markedly lower performance for our classifier compared to Uniref-90 and Uniref-100, and it is currently unclear whether the poor classification performance is due to the smaller size of the sequence dataset or the more stringent threshold on sequence identity. Surprisingly, the modified non-redundant sequence dataset^49^ from I-TASSER (PSSpred) gives a very high MCC value (0.457), even though it was released in 2014 and has not been continually updated as the other three. Thus, the DI-based classification performance depends on the sequence database in a nontrivial way and does not necessarily yield improved results for updated database versions.

**Figure 6.**
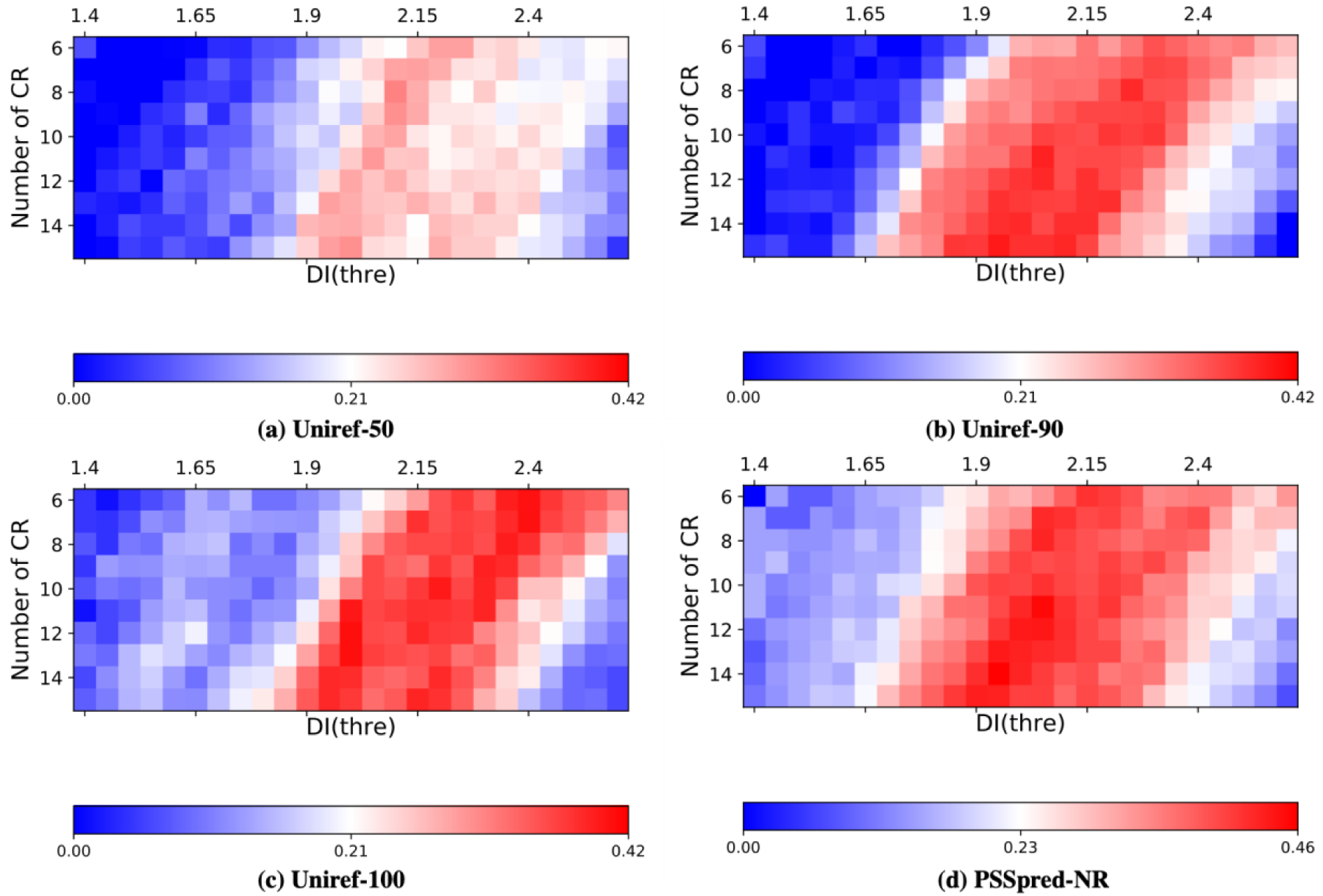
The different MCC values calculated based on: (a) 50% non-redundant sequence dataset, (b) 90% non-redundant sequence dataset, (c) 100% non-redundant sequence dataset and (d) PSSpred non-redundant sequence dataset.

## Conclusions

In this paper we described a diversity index-based classification model to predict metamorphic behavior in proteins solely based on the protein sequence. Our model was trained on a reference dataset consisting of 140 known monomorphic proteins and 201 known metamorphic proteins. Although the main purpose of SSP programs is to predict secondary structure, our results indicate that the “byproducts” of SSP, namely the alternate SS probabilities and the derived diversity index, can play a key role for predicting metamorphism in proteins. Among the four popular SSP programs, SPIDER2 has the overall best performance and robustness in classifying proteins as monomorphic vs. metamorphic. Further improvements in performance may be obtained by comparing the output of multiple SSP programs. All of these results indicate that the diversity index is a key variable to represent the metamorphic property of proteins. Because all four SSP programs give similar MCC values when used in classification to within ∼10%, we think further improvements in predicting protein metamorphism will require SSP methods that focus more on accurate quantification of uncertainty rather than yielding the best fit to experimental data.

## Author Contributions

N.C. conceived the presented idea and carried out the research. N.C., A.L. and L.-P.W. designed the research. N.C. and L.-P.W. wrote the manuscript. M.D. and A.L. provided important feedback on the research and manuscript including ideas for data analysis and the broader context of the research. All authors provided feedback and revisions on the manuscript.

## Acknowledgements

This research was supported by the U.S. Army Research Office Award #W911NF-17-1-0434. We would like to acknowledge Archana Chavan, Xuejun Yao, and Yudong Qiu for valuable discussions.

